# Diabetes and dementia incidence in Latin America; a 10/66 population-based cohort study

**DOI:** 10.1101/148155

**Authors:** Aquiles Salas, Daisy Acosta, Mariella Guerra, Yueqin Huang, Ivonne Z Jimenez-Velazquez, Juan J Llibre Rodriguez, Ana Luisa Sosa, Michael E Dewey, Ciro Gaona, Maëlenn M Guerchet, Loida Gonzalez, Zhaorui Liu, Jose A Luchsinger, A M Lopez Medina, Rosa M Salinas, Martin J Prince

## Abstract

**Background:** Diabetes prevalence is already high in middle income countries, particularly among older people. Current evidence on diabetes as a risk factor for dementia is limited to cohort studies in high income countries. Few studies carried out fasting glucose assessments to identify undiagnosed cases, and assess diabetes control. We aimed to determine the association between both diagnosed diabetes and total diabetes (including undiagnosed cases) and incident dementia, examining also the impact of glycaemic control on dementia risk.

**Methods:** Population-based cohort studies of those aged 65 years and over in sites in Cuba, Dominican Republic, Puerto Rico, Peru, Venezuela, and Mexico. Diagnosed diabetes was assessed through self-reported diagnosis, and undiagnosed diabetes through fasting blood samples (glucose >= 7mmol/L). Blood pressure, smoking, underactivity and waist circumference were assessed from questionnaires and physical examination. Incident 10/66 dementia (and subtypes), and mortality, were ascertained three to five years later.

**Results:** 12,297 interviews were completed at baseline, with 80-95% responding by site. The ‘at risk’ cohort comprised 10,945 dementia-free individuals, of whom 8,171 (75%) provided blood samples. Mean age varied from 72.0 to 75.1 years by site. Total diabetes prevalence was 43.5% in Puerto Rico, ranging from 11.5% to 27.0% in other sites. Most diabetes cases (50.2% to 68.4%) were not controlled (fasting glucose >7.0 mmol/L). 7,000 participants were followed up for 26,423 person-years with 659 incident dementia cases, and 905 dementia free deaths. Total diabetes was associated with incident 10/66 dementia (pooled meta-analysed adjusted sub-hazard ratio [pASHR] 1.25, 95% CI, 1.05-1.49, I^2^=48.6%), with a stronger association for uncontrolled (pASHR 1.47, 95% CI 1.19-1.81, I^2^=49.6%) than controlled cases (pASHR 1.29, 95% CI 0.95-1.74, I^2^=13.3%). Total diabetes was strongly associated with the incidence of vascular dementia (pASHR 2.25, 95% CI 1.24-4.08, I^2^=23.7%), but not Alzheimer’s Disease (pASHR 0.99, 95% CI 0.70-1.42, I^2^=49.0%).

**Conclusions:** Diabetes, particularly when poorly controlled, may increase dementia risk. There is considerable scope for improved detection and control of diabetes among older people in these settings, and hence an opportunity to carry out proof of concept prevention trials. Overlapping epidemics of these age dependent disorders will challenge poorly-resourced health systems in the future.

## 1 Introduction

Diabetes prevalence increases sharply with age, from 2.4% in those aged 20-39 years to 21.6% among those aged 65 years and over, and in recent years the largest increases have occurred in the oldest age groups [1]. Prevalence in emerging economies in Latin America and China is already similar to that in the USA [2–4]. The general trend, in both developed and developing countries, is towards an increasing prevalence, linked to increases in sedentary life-styles and obesity [5,6]. The high prevalence of diabetes, together with unrealized potential for prevention and treatment, makes it potentially one of the most important modifiable risk factors for dementia.

In a 2014 review of modifiable risk factors for dementia, limitations were noted in existing systematic reviews of longitudinal studies testing for a prospective association of diabetes with the onset of dementia [7]. These were only partly overlapping in content, did not distinguish between mid-life and late-life exposure to diabetes, or between ‘diagnosed diabetes’ and ‘total diabetes’ (including undiagnosed cases identified through fasting glucose or Oral Glucose Tolerance Tests), and included health record database linkage studies that are prone to ascertainment bias. A subsequent review shares these limitations, and focuses only on Alzheimer’s Disease (AD) [8]. The 2014 systematic review [7] merged studies from the two most recent reviews [9,10] adding any further eligible studies identified in the course of a new search. Any prospective or historical population-based cohort studies reporting on the association of diabetes with any dementia, Alzheimer’s Disease (AD) or vascular dementia (VaD) were included, but health record linkage studies were excluded other than for narrative description and discussion. Nineteen eligible studies were identified, and the meta-analyses were stratified by the stage in the life course at which the exposure had been ascertained (mid-life or late-life), and a sensitivity analysis was restricted to those studies in which undiagnosed as well as diagnosed diabetes had been ascertained. For late-life dementia, associations with incident AD (15 studies, pooled RR 1.40, 95% CI 1.22-1.61) were weaker than those with incident VaD (12 studies, pooled RR 2.39, 95% CI 1.92-2.98), with an intermediate effect size for any dementia (11 studies, pooled RR 1.50, 95% CI 1.33-1.70). For each outcome roughly half of the studies had included undiagnosed as well as diagnosed diabetes; the meta-analysed effect sizes were very similar in this subset of studies. No significant heterogeneity was noted for any of the meta-analyses (Higgins I^2^ =0.0%).

There are, nonetheless, important unanswered questions. All of the studies reviewed were conducted in high income countries, mainly in Europe and North America, with just one study from South Korea, and one from Japan. None of the studies was conducted in Latin America. In the Sacramento Area Latino Study on Aging, Mexican Americans aged 60 years and were followed up for 10 years, both treated diabetes (RR 2.05, 95% CI 1.41-2.97) and untreated diabetes (RR 1.55, 95% CI 0.93-2.58) were associated with increased risk of developing the composite outcome of either dementia or cognitive impairment. Recently published findings from the Mexico sites of the 10/66 Dementia Research Group (10/66 DRG) indicated that participants with type 2 diabetes mellitus had a near two-fold increased risk of developing dementia (RR 1.9; 95% CI 1.3-2.6) after three years of follow-up [11]. The impact of diabetes control is unclear with no studies comparing incident dementia risk between diagnosed and controlled and uncontrolled (whether diagnosed or undiagnosed) diabetics. Finally, the extent to which any association with diabetes may be mediated or confounded by other components of cardiovascular risk is uncertain, with few studies adjusting for a wider set of relevant risk factors.

Accordingly, we set out to study associations between diagnosed diabetes, total diabetes and the incidence of dementia in the 10/66 Dementia Research Group’s population-based cohort studies in Latin America where fasting blood samples had been taken at baseline, and assayed for plasma glucose. Our primary hypothesis was that total diabetes would be independently associated with an increased incidence of dementia. Secondary hypotheses were that a) there would be a linear association between glucose level and the risk of incident dementia, and b) that any increased risk of dementia would be concentrated among those with uncontrolled, as opposed to controlled diabetes. We planned further analyses to estimate associations of total diabetes with Alzheimer’s disease and vascular dementia subtypes. As noted, some but not all of these analyses have been previously reported for our Mexico sites [11], which are included here in revised form to maintain consistency with analysis protocols for other sites, for contextualisation and meta-analytical summary.

## Materials and methods

The 10/66 population-based study protocols for baseline and incidence waves are published in an open access journal [12], and the cohort profile has been described [13]. Relevant details are provided here. Fasting blood glucose level was assessed at baseline in seven sites in six countries (urban sites in Cuba, Dominican Republic, Puerto Rico, Peru and Venezuela, and urban and rural sites in Mexico). Self-report of diagnosed diabetes was also obtained in all of these sites. The one-phase population-based surveys were carried out of all residents aged 65 years and over in the geographically defined catchment areas between 2003 and 2007, other than in Puerto Rico (2007-2009)[12]. The target sample was 2000 for each country, and 3000 for Cuba. There were few differences in baseline characteristics of those who did and did not provide samples, and those that were statistically significant were generally of small effect [4]. Self-reports of diagnosed diabetes were not associated with giving blood samples in any site [4].

The baseline survey included clinical and informant interviews, and physical examination. Incidence waves were subsequently completed, with a mortality screen, between 2007 and 2011 (2011-2013 in Puerto Rico) aiming for 3-4 years follow-up in each site [14]. Assessments were identical to baseline protocols for dementia ascertainment, and similar in other respects. We revisited participants’ residences on up to five occasions. When no longer resident we sought information on their vital status and current residence, from other contacts recorded at baseline. Where participants had moved away, we sought to re-interview them, even outside the catchment area. If deceased, we recorded the date, and completed an informant verbal autopsy, including evidence of cognitive and functional decline suggestive of dementia onset between baseline assessment and death [15].

## Measures

The 10/66 population-based study interview covers dementia diagnosis, mental disorders, physical health, a physical examination and anthropometry, demographics, a risk factor questionnaire, disability, health service utilisation, and care arrangements [12]. Only relevant assessments are detailed here.

### Diabetes

Diagnosed diabetes was defined as a self-reported medical diagnosis of diabetes (answering ‘yes’ to the question “have you ever been told by a doctor that you have diabetes?”). Blood samples were taken early in the morning after an overnight fast, using a fluoride oxalate sample bottle for plasma glucose estimation [4]. Samples were transported, on ice, to local laboratories for processing and assay. Total diabetes was defined as a self-reported medical diagnosis (diagnosed diabetes), or a fasting glucose of > 7 mmol/l in the absence of a previous diagnosis (undiagnosed diabetes). Those with a fasting glucose of > 7 mmol/l were considered ‘uncontrolled’ regardless of diagnosis, while those with diagnosed diabetes below this threshold were considered ‘controlled’. Glycated haemoglobin (haemoglobin A1c – HbA1c) provides a summary indicator of glycaemic control over the previous two to three months, but this assay was only conducted in our Puerto Rico site.

### Confounders and other covariates

Age, sex, education level (none, did not complete primary, completed primary, secondary or tertiary), smoking history, and underactivity (“not at all” or “not very” physically active) were all assessed in the baseline questionnaire. Waist circumference was measured in centimetres using a flexible tape measure. Central obesity was defined as a waist circumference of more than 40 inches (101.6 centimetres) in men and of more than 35 inches (88.9 centimetres) in women [16]. Those with self-reported hypertension (“have you ever been told by a doctor that you have high blood pressure?”) and/ or a blood pressure measurement meeting WHO/ International Society of Hypertension criteria (systolic blood pressure >=140 mmHg and/ or diastolic blood pressure >= 90 mmHg) were considered to have hypertension.

### Dementia

10/66 dementia diagnosis was allocated to those scoring above a cutpoint of predicted probability for dementia, calculated using coefficients from a logistic regression equation developed, calibrated and validated cross-culturally in the 25 centre 10/66 pilot study [17], applied to outputs from a) a structured clinical interview, the Geriatric Mental State [18], b) two cognitive tests; the Community Screening Instrument for Dementia (CSI-D) COGSCORE [19] and the modified CERAD 10 word list learning task with delayed recall [20], and c) informant reports of cognitive and functional decline from the CSI-D RELSCORE [19]. The prevalence [21] and incidence [15] of 10/66 dementia in the current cohorts have been reported. The criterion, concurrent and predictive validity of the 10/66 diagnosis were superior to that of the DSM-IV dementia diagnostic criterion in subsequent evaluations [21–24].

For those who died between baseline and follow-up we diagnosed ‘probable incident dementia’ by applying three criteria:

1. A score of more than two points on the RELSCORE, from the post-mortem informant interview, with endorsement of either ‘deterioration in memory’ or ‘a general deterioration in mental functioning’ or both, and

2. an increase in RELSCORE of more than two points from baseline, and

3. the onset of these signs noted more than six months prior to death.

In the baseline survey, the first criterion would have detected those with either DSM-IV or 10/66 dementia with 73% sensitivity and 92% specificity [15].

For those who survived to the follow-up interview, information from cognitive test scores, neurological examination and History and Aetiology Schedule (Dementia Diagnosis and Subtype) was used to allocate subtype diagnoses using a computerized algorithm to apply relevant criteria. Subtype allocation was therefore not possible for ‘probable incident dementia’ cases identified from post-mortem informant interview. We focused upon the two commonest dementia subtypes:

a) pure or mixed case of possible or probable AD according to National Institute of Neurological and Communicative Disorders and Stroke and the Alzheimer’s Disease and Related Disorders Association criteria (NINCDS/ADRDA) criteria [25]

b) pure or mixed cases of possible Vascular Dementia (VaD) according to National Institute of Neurological Disorders and Stroke and Association Internationale pour la Recherché et l’Enseignement en Neurosciences criteria (NINDS-AIREN) criteria [26].

## Analyses

We used release 2.0 of the 10/66 dementia incidence data archive (October 2015), and STATA version 11 for all analyses.

For each site

1. we describe the baseline characteristics of the at risk cohort (dementia-free participants with blood tests) and their status at follow-up.

2. we modelled the effect of diabetes exposures on the incidence of dementia (10/66 dementia or ‘probable dementia among those who had died) using a competing-risks regression derived from Fine and Gray’s proportional subhazards model [27] (Stata stcrreg command), based on a cumulative incidence function, indicating the probability of failure (dementia onset) before a given time, acknowledging the possibility of a competing event (dementia-free death). Time to death was the time from baseline interview to the exact date of death. Time to dementia onset (which could not be ascertained precisely) was the midpoint between baseline and follow-up interview. Competing risks regression keeps those who experience competing events at risk so that they can be counted as having no chance of failing. We report adjusted sub-hazard ratios (ASHR) with robust 95% confidence intervals adjusted for household clustering. For the primary analysis, all models were adjusted for age, sex, and education, and then extended to control also for hypertension, obesity, smoking, and underactivity. All models were estimated separately for each site, and the results combined using a fixed effects meta-analysis. Higgins I^2^ estimates the proportion of between-site variability in the estimates accounted for by heterogeneity, as opposed to sampling error; up to 40% heterogeneity is conventionally considered negligible, while up to 60% reflects moderate heterogeneity [28].

We classified diabetes exposure in three ways; for the primary analysis, total diabetes (whether controlled or uncontrolled) versus no diabetes; total diabetes stratified by control versus no diabetes; and the linear effect of serum glucose level (per mmol/l). To assess possible bias arising from reliance only on self-reported diabetes diagnoses, we estimated the effect of this exposure on the incidence of 10/66 dementia, for those who had provided blood samples, the reference group being all those with no diagnosed diabetes. We also carried out secondary analyses exploring the association of total diabetes with the specific sub-types of Alzheimer’s disease (AD) and vascular dementia (VaD); those with other subtypes of dementia, or for whom a subtype could not be allocated were excluded from these analyses.

The study protocol and the consent procedures were approved by the King’s College London research ethics committee and in all countries where the research was carried out: 1- The Memory Institute and Related Disorders (IMEDER) Ethics Committee (Peru); 2-Finlay Albarran Medical Faculty of Havana Medical University Ethical Committee (Cuba); 3-Hospital Universitario de Caracas Ethics Committee (Venezuela); 4- Consejo Nacional de Bioética y Salud (CONABIOS, Dominican Republic); 5- Instituto Nacional de Neurología y Neurocirugía Ethics Committee (Mexico); 6- University of Puerto Rico, Medical Sciences Campus Institutional Review Board (IRB).

## Results

### Cohort characteristics

For the seven sites in six countries where blood samples were obtained, 12,297 interviews were completed at baseline (see Table 1). The ‘at risk’ cohort comprised 10,945 dementia-free participants of whom 8,171 (75%) had provided blood samples with valid serum glucose levels recorded. Mean age at baseline for these participants ranged from 72.0 to 75.1 years. Educational levels were lowest in rural Mexico (84% not completing primary education), and Dominican Republic (70%), and highest in Puerto Rico (20%), and Cuba (23%). The prevalence of total diabetes was exceptionally high in Puerto Rico (43.5%), and ranged from 11.5% to 27.0% in other sites. In all sites, most diabetes cases were uncontrolled (fasting glucose >7.0 mmol/ L), the proportion varying from 50.2% to 68.4% by site. Across the cohort, the prevalence of hypertension was 72.7% (range by site 53.8% to 77.9%), obesity 46.8% (39.0% to 57.1%), ever smoker 37.5% (26.1% to 48.0%), and underactivity 30.6% (26.9% to 34.5%).

**Table 1.** Cohort description Cuba Dominican

|  | Cuba | Dominican Republic | Puerto Rico | Peru (urban) | Venezuela | Mexico (urban) | Mexico (rural) | All sites |
| --- | --- | --- | --- | --- | --- | --- | --- | --- |
| Interviewed at baseline (response %) | 2928 (94%) | 2011 (95%) | 2009 (93%) | 1381 (80%) | 1965 (80%) | 1003 (84%) | 1000 (86%) | 12297 |
| Interviewed at baseline with fasting glucose | 2355 (80.4%) | 1483 (73.7%) | 1586 (78.9%) | 770 (55.8%) | 1284 (65.3%) | 822 (82.0%) | 895 (89.5%) | 9195 (74.8%) |
| Dementia free cohort | 2517 | 1769 | 1765 | 1251 | 1820 | 910 | 913 | 10945 |
| Dementia free cohort with fasting glucose | 2012 (79.9%) | 1278 (72.2%) | 1435 (81.3%) | 689 (55.1%) | 1193 (65.5%) | 748 (82.2%) | 816 (89.4%) | 8171 (74.7%) |
| Follow-up tracing |  |  |  |  |  |  |  |  |
| Interviewed | 1522 (75.6%) | 835 (65.3%) | 1082 (75.4%) | 531 (77.1%) | 852 (71.4%) | 599 (80.1%) | 592 (72.5%) | 6013 (73.6%) |
| Deceased | 350 (17.4%) | 249 (19.5%) | 130 (9.1%) | 29 (4.2%) | 93 (7.8%) | 57 (7.6%) | 79 (9.7%) | 987 (12.1%) |
| Refused | 10 (0.5%) | 20 (1.6%) | 3 (0.2%) | 89 (12.9%) | 111 (9.3%) | 38 (5.1%) | 91 (11.2%) | 362 (4.4%) |
| Lost | 130 (6.4%) | 174 (13.6%) | 220 (15.3%) | 40 (5.8%) | 137 (11.5%) | 54 (7.2%) | 54 (6.7%) | 809 (9.9%) |
| Person years of follow-up | 7562 | 4759 | 4970 | 1682 | 3725 | 1886 | 1838 | 26423 |
| Follow-up outcome |  |  |  |  |  |  |  |  |
| Incident dementia cases | 159 (8.7%) | 121 (11.2%) | 129 (10.8%) | 23 (4.1%) | 113 (12.0%) | 38 (5.8%) | 76 (11.3%) | 659 (9.5%) |
| Competing risk (dementia-free death) | 340 (18.5%) | 221 (20.4%) | 107 (8.9%) | 27 (4.8%) | 82 (8.7%) | 54 (8.2%) | 74 (11.0%) | 905 (13.0%) |
| Censored | 1339 (72.9%) | 742 (68.5%) | 964 (80.3%) | 509 (91.1%) | 750 (79.4%) | 564 (86.0%) | 521 (77.6%) | 5389 (77.5%) |
| Cohort characteristics at baseline |  |  |  |  |  |  |  |  |
| Age, mean (SD) | 74.3 (6.6) | 74.3 (6.9) | 75.1 (6.7) | 73.8 (5.6) | 72.0 (6.4) | 73.6 (6.0) | 73.6 (6.3) | 73.9 (6.6) |
| Sex (female %) | 1319 (65.6%) | 871 (68.2%) | 970 (67.8%) | 444 (64.4%) | 768 (64.4%) | 498 (66.6%) | 493 (60.4%) | 5363 (65.7%) |
| Education (did not complete primary, %) | 452 (22.5%) | 897 (70.4%) | 282 (19.7%) | 47 (6.9%) | 344 (29.0%) | 400 (53.7%) | 682 (83.6%) | 3104 (38.1%) |
| Glucose (mmol/L), mean (SD) | 5.6 (2.1) | 5.3 (2.0) | 6.6 (2.8) | 5.6 (1.4) | 5.5 (2.5) | 5.6 (2.6) | 6.0 (2.6) | 5.8 (2.4) |
| Total diabetes (%) | 488 (24.4%) | 229 (17.9%) | 624 (43.5%) | 79 (11.5%) | 235 (21.0%) | 202 (27.0%) | 196 (24.0%) | 2053 (25.4%) |
| Controlled diabetes | 231 (11.6%) | 114 (8.9%) | 207 (14.4%) | 29 (4.2%) | 102 (9.1%) | 81 (10.8%) | 62 (7.6%) | 826 (10.2%) |
| Uncontrolled diabetes | 257 (12.9%) | 115 (9.0%) | 417 (29.1%) | 50 (7.3%) | 133 (11.9%) | 121 (16.2%) | 134 (16.4%) | 1227 (15.2%) |
| Proportion of diabetes cases controlled | 47.3% | 49.8% | 33.2% | 36.7% | 43.4% | 40.0% | 31.6% | 40.2% |
| Hypertension | 1516 (75.5%) | 995 (77.9%) | 1113 (79.6%) | 370 (53.8%) | 879 (78.2%) | 521 (69.7%) | 484 (56.9%) | 5858 (72.7%) |
| Obesity | 782 (39.0%) | 625 (49.2%) | 714 (54.9%) | 287 (42.2%) | 487 (48.8%) | 425 (57.1%) | 337 (41.6%) | 3657 (46.8%) |
| Ever smoked | 909 (45.3%) | 613 (48.0%) | 373 (26.1%) | 155 (22.5%) | 518 (43.9%) | 261 (34.9%) | 224 (27.5%) | 2053 (37.5%) |
| Underactivity | 539 (26.9%) | 436 (34.2%) | 417 (29.2%) | 197 (27.2%) | 410 (34.5%) | 254 (34.1%) | 243 (30.0%) | 2486 (30.6%) |

From the ‘at risk’ cohort, 362 participants (4.4%) refused, and 809 (9.9%) were lost to follow-up; 7,000 participants with baseline fasting glucose blood counts (85.7%) were followed up for 26,423 person years (Table 1). In total, 659 incident cases of dementia were identified (9.5%), while a competing risk of dementia free death was recorded for 905 (13.0%); 5,389 (77.5%) participants were dementia-free at follow-up. Dementia outcomes could not be obtained for 47 participants due to incomplete follow-up interviews, and these were excluded from subsequent analyses. Of the 577 incident cases of 10/66 dementia, 199 were allocated to the AD subtype, and 59 to the VaD subtype; 290 did not fully meet criteria for any subtype, and 35 met criteria for other subtypes.

## Primary analysis - total diabetes and the incidence of dementia

The pooled meta-analysed adjusted sub-hazard ratio (ASHR) for the association of total diabetes with incident 10/66 dementia, controlling for age, sex and education was 1.25 (95% CI, 1.05-1.49) with moderate heterogeneity among sites (I^2^=48.6%) (Table 2). The association was most prominent in urban Mexico and rural Mexico, with an association of borderline statistical significance in Puerto Rico. After controlling further for hypertension, obesity, smoking, and underactivity, the association was generally stronger, with less heterogeneity of effect among sites (pooled ASHR 1.36, 95% CI 1.12-1.63, I^2^=39.5%).

**Table 2.**

| Exposure | Diagnosed diabetes | Total diabetes | Total diabetes | Total diabetes stratified by control |  | Glucose level |
| --- | --- | --- | --- | --- | --- | --- |
| Non-exposed (reference group) | No diagnosed diabetes | No diabetes | No diabetes | No diabetes |  | Per mmol/ L |
|  | Adjusted ASHR (base model<sup>1</sup>) | Adjusted ASHR (base model<sup>1</sup>) | Adjusted ASHR (extended model<sup>2</sup>) | Adjusted ASHR (base model<sup>1</sup>) |  | Adjusted ASHR (base model<sup>1</sup>) |
| Cuba | 0.80 (0.53-1.23) | 0.81 (0.56-1.19) | 0.87 (0.59-1.28) | Controlled | 0.61 (0.34-1.09) | 0.99 (0.92-1.06) |
|  |  |  |  | Uncontrolled | 1.00 (0.63-1.58) |  |
| Dominican Republic | 0.95 (0.56-1.60) | 1.14 (0.72-1.83) | 1.33 (0.82-2.16) | Controlled | 0.48 (0.19-1.16) | 1.11 (1.04-1.20) |
|  |  |  |  | Uncontrolled | 1.96 (1.15-3.33) |  |
| Puerto Rico | 1.30 (0.91 -1.86) | 1.38 (0.99-1.94) | 1.59 (1.09-2.31) | Controlled | 1.37 (0.84-2.23) | 1.06 (1.00-1.13) |
|  |  |  |  | Uncontrolled | 1.39 (0.95-2.03) |  |
| Peru, urban | 0.57 (0.07-4.30) | 0.46 (0.06-3.28) | 0.42 (0.06-3.26) | Controlled | 1.11 (0.15-8.40) | 0.79 (0.59-1.07) |
|  |  |  |  | Uncontrolled | No incident dementia cases |  |
| Venezuela | 1.45 (0.89-2.36) | 1.24 (0.78-1.96) | 1.36 (0.83-2.21) | Controlled | 1.67 (0.94-2.97) | 0.99 (0.90-1.08) |
|  |  |  |  | Uncontrolled | 0.95 (0.49-1.82) |  |
| Mexico, urban | 2.59 (1.35-4.94) | 2.23 (1.16-4.27) | 2.01 (1.05-3.85) | Controlled | 1.34 (0.45-3.93) | 1.16 (1.06-1.26) |
|  |  |  |  | Uncontrolled | 2.80 (1.39-5.66) |  |
| Mexico, rural | 1.36 (0.78-2.39) | 1.81 (1.10-2.99) | 1.88 (1.11-3.16) | Controlled | 1.49 (0.63-3.55) | 1.07 (1.01-1.14) |
|  |  |  |  | Uncontrolled | 1.95 (1.12-3.40) |  |
| **Pooled estimates** | **1.22 (1.00-1.47)** | **1.25 (1.05-1.49)** | **1.36 (1.12-1.63)** | **Controlled** | **1.29 (0.95-1.74)** | **1.06 (1.03-1.09)** |
|  |  |  |  | **Uncontrolled** | **1.47 (1.19-1.81)** |  |
| **Higgins I<sup>2</sup>** | **46% (0-77%)** | **49% (0-78%)** | **40% (0-75%)** | **Controlled** | **13% (0-78%)** | **61% (10-83%)** |
|  |  |  |  | **Uncontrolled** | **50% (0-80%)** |  |

1. controlling for age, sex and education level
2. controlling for age, sex, education level, hypertension, obesity, underactivity and smoking

## Secondary analyses

The association between self-reported diagnosed diabetes and the incidence of 10/66 dementia was similar to the association with total diabetes in all sites, and when pooled meta-analytically (pooled ASHR 1.22, 95% CI 1.00-1.47, I^2^=46.3%).

Stratifying the total diabetes exposure according to glycaemic control indicated that risk for incident dementia may be concentrated among those with uncontrolled diabetes (pooled ASHR 1.47, 95% CI 1.19-1.81, I^2^=49.6%), rather than controlled diabetes (pooled ASHR 1.29, 95% CI 0.95-1.74, I^2^=13.3%). The linear association of serum glucose level (per mmol/ L) with incident 10/66 dementia was strongly significant when pooled across sites, but with substantial heterogeneity of effect (pooled ASHR 1.06, 95% CI 1.03-1.09, I^2^=60.9%). In the Puerto Rico site, where HbA1c concentrations were estimated, this indicator of longer-term control was associated with the incidence of 10/66 dementia in the cohort as a whole (ASHR per SD 1.22, 95% CI 1.02-1.46), and among diabetes cases (ASHR per SD 1.26, 95% CI 1.04-1.54)

There was no association between total diabetes and the incidence of AD (pooled ASHR 0.99, 95% CI 0.70-1.42, I^2^=49%, 95% CI 0-80%). However, total diabetes was strongly associated with the incidence of vascular dementia, with negligible heterogeneity among sites (pooled ASHR 2.25, 95% CI 1.24-4.08, I^2^=24%, 95% CI 0-68%).

## Discussion

### Principal findings

In a large population-based cohort study in rural and urban sites in Latin America, we demonstrated an association between total diabetes (diagnosed and undiagnosed cases) and the incidence of any dementia, with only low to moderate heterogeneity in effect sizes among sites. The increased risk was somewhat concentrated among those with uncontrolled diabetes. For the subset of incident cases among whom dementia subtype could be allocated, there was a strong association between total diabetes and the onset of vascular dementia, and no association with the onset of AD. The association between diagnosed diabetes and the incidence of any dementia was similar, but slightly smaller than that for total diabetes.

### Strengths and weaknesses of the study

Strengths of this study are that associations have been assessed longitudinally, in large population-based dementia-free cohorts, encompassing rural and urban catchment area sites in Latin America and the Caribbean. Given modest heterogeneity, we were able to use meta-analytical techniques to pool effect sizes and increase the precision of our estimates. Fasting glucose assays enabled us to identify undiagnosed as well as diagnosed cases of diabetes, and to assess the impact of glycaemic control on dementia risk. We were able to control for the potential confounding effects of an extensive set of known covariates of diabetes from these settings [4], including other cardiovascular risk factors.

We acknowledge some limitations. First there will have been some misclassification of the self-reported exposure of diagnosed diabetes, which, even given the prospective design, may have been differential with respect to the outcome, with those in the process of developing dementia perhaps being less likely to recall or report a valid diagnosis. The effect of this, and any random misclassification, would have been towards an underestimation of any genuine association. There will also have been some measurement error in fasting glucose estimations. Although all samples were stored in fluoride oxalate, and transported on ice to the local laboratory, time to processing will have varied, and longer delays will have led to underestimation of true plasma levels due to glycolysis. This effect can be presumed to be random with respect to the outcome, leading again to an underestimation of the true risk effect. Diabetes diagnostic precision would have been enhanced if oral glucose tolerance tests were carried out, but this was impractical in the community-setting of these epidemiological studies. Evaluation of glycaemic control was hampered by the lack, other than in Puerto Rico, of glycated haemoglobin assays, which would have given a better summary of recent control than a one-off fasting glucose.

### Contextualisation with other research

Our findings are broadly consistent with those previously reported from a range of high income countries (see introduction and [7]), although the effect size for the association of any dementia with total diabetes is marginally weaker, and we found no association between total diabetes and the incidence of AD. Consistent with the findings from the earlier meta-analysis [7], estimation of the association based upon self-reported diagnosis only, misclassifying undiagnosed cases, did not seem to lead to any consistent or substantial bias.

An update of the earlier meta-analysis [7], to which the seven 10/66 sites providing data on total diabetes contribute 30% of the weight, suggests a pooled relative risk of 1.46 (95% CI 1.32-1.61) for the outcome of any dementia, with negligible heterogeneity (I^2^ =11%, 95% CI). For AD the updated pooled relative risk is 1.34, 95% CI 1.18 to 1.52, I^2^=21%, 95% CI 0-54% (with the 10/66 cohorts contributing 13% of the weight) and for VaD 2.38, 95% CI 1.94-2.92, I^2^=0.0%, 95% CI 0-50% (12% of the weight). Our study therefore extends existing evidence of an association with dementia, to middle income countries in Latin America where the prevalence of diabetes is high and control inadequate [4].

None of the seven previous population-based studies that have used plasma glucose assessment or oral glucose tolerance tests in addition to self-reported diagnoses to ascertain diabetes cases at baseline has reported on subsequent dementia risk stratified by diabetes control [29–35]. Our finding of an apparent concentration of risk among uncontrolled cases is therefore original, and the case for the salience of diabetes control is further strengthened by the observation, in the Puerto Rico site, of an association between glycated haemoglobin level and dementia risk. Our results are consistent with findings from diabetes cohorts that worse glycaemic control predicts cognitive decline [36], and that diabetes complications (indicators of poor glycaemic control) are associated with an increased risk of dementia [37,38].

### Possible mechanisms

It remains possible that processes or genetic predispositions that underlie both diabetes and AD [39], could explain a link that is not causal. It is also possible that diabetes decreases brain resilience, but does not directly cause dementia or AD. There are, however, several possible mechanisms by which diabetes may increase the risk of dementia and AD. Insulin resistance is an antecedent and correlate of diabetes, and is considered to be a central mechanism in the metabolic syndrome, but few studies have demonstrated an association with dementia [40,41]. The association of the metabolic syndrome with dementia is inconsistent, with some studies indicating a specific association with the diabetes component alone [29,42]. Cardiovascular and cerebrovascular disease are supported as important mediating mechanisms by our finding, consistent with evidence from previous studies [7], that diabetes is a much stronger risk factor for vascular dementia than for AD. In a large population-based autopsy series from Finland, diabetes in late-life was positively associated with cerebral infarcts, but not with either β-amyloid or neurofibrillary tangles [43]. However, diabetes may also directly affect AD neuropathological processes [44]. Brain imaging studies of cognitively normal individuals indicate that insulin resistance [45] and diabetes [46] are related to a regional profile of reduced brain metabolism consistent with AD. In a study from South Korea anti-amyloid β antibodies were elevated in persons with diabetes, possibly mediated by dyslipidaemia [47]. Advanced glycation endproducts (AGEs) are elevated in diabetes, where they are strongly implicated in end-organ damage. AGEs are also elevated in AD brains and stimulate beta-amyloid production [48]. In rat experimental models, treatment with AGEs can induce tau hyperphosphorylation and impair synapse and memory through upregulation of the AGEs receptor (RAGE) activating glycogen synthase kinase-3 (GSK-3), changes that are then reversed by blocking the RAGE/ GSK-3 pathway [48].

### Implications for dementia risk reduction

Our evidence confirms a consistent association between diabetes in late-life and the subsequent onset of dementia. This is in contrast to the pattern observed for hypertension, obesity and dyslipidaemia, where increased risk is only apparent for midlife exposures [7]. If causal, the association may have important implications for dementia risk reduction in later-life. However, it is unclear whether prevention or more effective treatment of diabetes can prevent dementia [49], since, in contrast to other cardiovascular risk factors, few randomized controlled trials have been conducted. Two treatment trials give conflicting evidence. In the ACCORD MIND trial people aged 55-80 years with poorly controlled diabetes and elevated cardiovascular risk who were randomized to tighter than normal glycaemic control had similar cognitive function after 40 months to the control arm aiming for normal glycaemic control, although neuroimaging suggested less brain atrophy [50]. In another clinical trial among diabetics aged 55 years and over, those randomized to a telemedicine intervention that improved glycaemic control experienced less global cognitive decline, an effect that seemed to be mediated by changes in HbA1c [51]. Two recent European trials of multidomain cardiovascular risk reduction interventions to prevent cognitive decline and dementia also give mixed findings [52,53]. In the Finnish Geriatric Intervention Study to Prevent Cognitive Impairment and Disability (FINGER) trial, participants, aged 60-77 years with elevated dementia risk scores, randomized to a 2 year intensive multidomain intervention (diet, exercise, cognitive training, vascular risk monitoring) showed more cognitive improvement than the those in the control group (general health advice) [52]. In contrast, in the Dutch preDIVA primary care cluster-randomised controlled trial of a 6-year nurse-led, multidomain cardiovascular intervention versus usual care there was no difference in dementia incidence between the two groups) [53]. These trials unfortunately contribute little to our understanding of the specific role of diabetes prevention and control, since fasting glucose levels did not differ between arms over the follow-up period in either trial, although there were greater reductions in body mass index in the intervention arm in FINGER [52], and better control of hypertension in preDIVA [53]. The preDIVA investigators commented that the high quality of ‘usual care’ in the Netherlands may have precluded demonstration of an intervention effect [53].

While there is minimal available evidence on the long latency association between diabetes in mid-life and the onset of dementia in late life [7], three health service register linkage studies, conducted in USA [54], Korea [55] and Finland [56] all support an association between midlife diabetes and dementia risk, which may be even stronger than for diabetes diagnosed in late-life. Therefore, efforts to optimize glycaemic control need to start early and should be maintained lifelong. However, hypoglycaemic attacks, an unintended consequence of tight glycaemic control, are a concern, particularly in older diabetes patients. Rapid improvement in glycaemic control (falling HbA1c levels, from a high baseline level) seems to be associated with worse cognitive outcomes than either stable good or bad control [57], and hypoglycaemic attacks strongly predict the onset of dementia [58]. Evidence would therefore support a cautious approach to optimizing glycaemic control in older diabetics, avoiding hypoglycaemia where possible. Hypoglycaemia may be a consequence as well as a cause of cognitive impairment [58], since the onset of cognitive impairment and dementia greatly complicate diabetes self-management and treatment [59]. American Diabetic Association Standards of Medical Care in Diabetes recommend a less stringent glycaemic control target (HbA1c<8.0%) for those with a severe hypoglycaemia history, limited life expectancy, advanced vascular complications, extensive comorbidities, and in long-term diabetes where targets are difficult to attain [60].

### Conclusions

Improved understanding of causal mechanisms may help to orientate future diabetes treatment and prevention strategies towards the prevention of dementia. In settings such as those included in this study, where management of diabetes is sub-optimal, trials of community and health system strengthening interventions to improve diabetes detection and control among older people are indicated. Such trials could usefully include cognitive decline and dementia as additional outcomes, since there may be more scope than in high income country health systems to demonstrate proof of concept of prevention potential. Regions undergoing rapid population ageing and economic development are experiencing rising, and overlapping epidemics of diabetes and dementia. It seems that comorbidity between these conditions is determined by more than their common age dependency. Nevertheless, population ageing will drive sharp increases in the numbers of older adults living with diabetes and comorbid cognitive impairment. Their impaired capacity for adherence to treatment and self-care, and less favourable outcomes will pose a challenge for healthcare systems worldwide, particularly those in less-resourced settings.

## Acknowledgements

The 10/66 Dementia Research Group’s research has been funded by the Wellcome Trust Health Consequences of Population Change Programme (GR066133 – Prevalence phase in Cuba and Brazil; GR080002- Incidence phase in Peru, Mexico, Cuba, Dominican Republic, Venezuela and China), the World Health Organisation (India, Dominican Republic and China), the US Alzheimer’s Association (IIRG – 04 – 1286 - Peru, Mexico and Argentina), and FONDACIT (Venezuela). The analysis reported here was carried out with funding support from the European Research Council (ERC-2013-ADG 340755 LIFE2YEARS1066). The Rockefeller Foundation supported our dissemination meeting at their Bellagio Centre.

